# Spatio-temporal dynamics and drivers of Highly Pathogenic Avian Influenza H5N1 in Chile

**DOI:** 10.1101/2023.04.24.538139

**Authors:** Claudio Azat, Mario Alvarado-Rybak, José Fernando Aguilera, Julio A. Benavides

## Abstract

Highly pathogenic avian influenza A H5N1 clade 2.3.4.4b (hereafter H5N1) is causing vast impacts on biodiversity and poultry around the globe. In Chile it was first reported on December 7^th^, 2022, in a pelican (*Pelecanus thagus*) found dead in the northern city of Arica. In the following months, lethal H5N1 cases were reported in a wide range of wild bird species, marine mammals, backyard and industrial poultry, and in a human. Despite its high impact and spread, it is not well-known what environmental factors are associated with outbreaks. This study describes the spatio-temporal patterns of the current epizootic of H5N1 in Chile and test ecological and anthropogenic drivers that could be associated with outbreak occurrence. We used H5N1 cases reported by the Chilean national animal health authority to the World Animal Health Information System (WAHIS) from December 9^th^, 2022, to March 3^rd^, 2023. These included bird cases confirmed through avian influenza specific real-time PCR assay (qPCR), obtained from passive and active surveillance. Data was analyzed to detect the presence of H5N1 clusters under space-time permutation probability modelling, H5N1 association between distance and days since first outbreak through linear regression, and correlation between H5N1 presence with a range of ecological and anthropogenic variables by general linear modelling. From the 197 H5N1 identified outbreaks, involving 478 individual cases among wild and domestic birds, a wave-like steady spread of H5N1 from north to south was identified, that can help predict hotspots of outbreak risk and establish targeted preventive measures. For instance, 14 statistically significant clusters were identified, with the largest located in central Chile (18-29 km in radius) where poultry production is concentrated. Also, one of the clusters was identified in Tocopilla, location where the H5N1 human case occurred time later. In addition, the presence of H5N1 outbreaks was positively correlated with bird richness, human footprint, precipitation of the wettest month, minimum temperature of the coldest month, and mean diurnal temperature. In contrast, presence of H5N1 was negatively correlated to distance to the closest urban center, precipitation seasonality and isothermality. Preventive actions based on our modeling approach include developing wildlife surveillance diagnostic capabilities in Chilean regions concentrating outbreaks. It is urgent that scientists, the poultry sector, local communities and national health authorities co-design and implement science-based measures from a One Health perspective to avoid further H5N1 spillover from wildlife to domestic animals and humans, including rapid removal and proper disposal of wild dead animals, and the closure of public areas (i.e., beaches) reporting high wildlife mortalities.

## INTRODUCTION

The current panzootic of Highly Pathogenic Avian Influenza A subtype H5N1 clade 2.3.4.4b (hereafter H5N1) is causing large mortalities of wild populations of birds and mammals worldwide (Gamarra-Toledo et al. 2023a, Leguia et al. 2023, Nemeth et al. 2023). In addition, >200 million domestic poultry have died or been culled as a control management (World Organization of Animal Health 2023), threatening food security of low- and middle-income countries. First described in the Netherlands in October 2020, the novel H5N1 virus bearing the clade 2.3.4.4b HA gene (Lewis et al. 2021) originated from reassorted avian influenza H1N1, H3N8 and H5N8 viruses (Lewis et al. 2021, Pohlmann et al. 2022, Shi et al. 2023). This novel H5N1 virus has spread rapidly through major avian migratory pathways to every continent except Australia and Antarctica so far (Caliendo et al. 2021, Bevins et al. 2022, Shi et al. 2023). Although it has major impacts on wild and domestic birds, the unusually high and growing number of lethal cases among marine and terrestrial mammals, especially in Europe and the Americas, are a hallmark of the current emergency (Bordes et al. 2023, Gamarra-Toledo et al. 2023b, Puryear et al. 2023). Unprecedented mammal-to-mammal transmission has been increasingly suspected. For instance, Agüero et al. (2023) described the mortality of several thousand farmed American minks (*Neovison vison*) in northern Spain following a rapid pen to pen pattern of spread. But also, spillover to people, with 10 cases of H5N1 human infections reported since January 2022, two of them fatal, fuel fears of the emergence of a new global pandemic of unsuspected consequences (Kupferschmidt 2023, Sidik 2023).

In Chile, the Agriculture and Livestock Service (SAG) reported the first case of H5N1 on December 7^th^, 2022, in a Peruvian pelican (*Pelecanus thagus*) found dead in the northern coastal city of Arica, confirmed by a specific reverse-transcriptase real-time PCR assay (hereafter qPCR; Heine et al. 2007). This occurred few weeks after Peru reported the mass mortality of >22,000 marine birds (mainly *P. thagus*) presumably due to avian influenza (Gamarra-Toledo et al. 2023a). Two months later, Chile reported the first case in backyard poultry in the coastal town of Chañaral de Aceituno, followed by the first confirmed cases in marine mammals in a South American sea lion (*Otaria flavescens*) and a marine otter (*Lontra felina*), also in the North. Subsequently, the first outbreak in industrial poultry in Central Chile was reported in March 2023 (in the city of Rancagua), which concentrates the country’s poultry production. More recently, on March 29^th^ the Ministry of Health reported Chile’s first case of H5N1 in a 53-year-old man from the coastal city of Tocopilla in northern Chile, raising the possibility of this panzootic becoming a serious threat to human health.

Despite multiple non-pathogenic and highly pathogenic avian influenza viruses circulating across the globe for decades, and previous pandemics including H1N1 Spanish flu (1918) and H1N1 swine flu (2009), there is still a large lack of understanding on how the novel H5N1 virus spreads, particularly given its high mortality and diverse range of affected hosts. A better understanding of the ecology of H5N1 is urgently needed to inform effective strategies for disease surveillance and control in Chile and elsewhere. This study describes the spatio-temporal patterns of the current epizootic of H5N1 in Chile and test ecological and anthropogenic drivers that could be associated with outbreak occurrence, to help predicting and preventing areas of high risk for future outbreaks in wildlife, domestic poultry, and potential cases in humans.

## METHODS

### Data collection

We used cases of H5N1 in wild and domestic birds reported from December 9^th^, 2022, to March 3^rd^ 2023 by the Chilean national animal health authority (SAG) to the World Animal Health Information System (WAHIS) platform (WAHO 2023). These were confirmed cases through qPCR targeting the avian influenza M gene, obtained from passive and active surveillance, including samples of live and dead wild birds, backyard production birds and industrial poultry. Sampling typically consisted of oral, tracheal, or cloacal swabs, but also included tissues (Ariyama et al. 2023). From each record we obtained species, date, geographical coordinates, condition (death or alive), and if animals were killed and disposed. An outbreak was considered as one entered line on the WAHIS system, and the total number of positive/suspected case was considered in the number of cases per outbreak.

### H5N1 spatio-temporal cluster analysis

Visualization of H5N1 outbreak locations was generated using QuantumGIS v.3.16.11 (QGIS Development Team 2023) and projected for analysis using WGS 1984 datum as a coordinate system. Each sampled site was considered as a H5N1 outbreak, if at least one individual swab sample tested positive, according to the results of the specific qPCR. First, spatial distribution was characterized by the global Moran’s I index (Carpenter 2001), to identify spatial autocorrelation in the country. Then, we used scan statistic (Kulldorff 1997) to detect any cluster of H5N1 outbreaks under space-time permutation probability modelling. The model was run using H5N1 locations under the null hypothesis that outbreaks were randomly distributed across the country. The model was set to scan for areas with high H5N1 positives numbers to test for clusters with a spatial occurrence higher than that outside the cluster. The model allows the detection of the most “unusual” excess of observed H5N1 outbreaks and therefore provides georeferenced high-risk areas of H5N1 occurrence. P-value (P<0.05) were obtained using Monte Carlo simulation by generating 999 replications of the data set and analysis was performed using the software SatScan v.10.1 (Kulldorff 2009).

### Ecological and anthropogenic H5N1 drivers

We first tested the association between distance and days since the first outbreak through a linear regression model. We also tested the correlation of H5N1 presence/absence of outbreaks in birds with several ecological and anthropogenic variables that could be related to disease occurrence or reporting. For this purpose, we built a global model including 15 out of the 19 bioclimatic variables from WorldClim (Fick and Hijmans 2017), excluding variable 8 (mean temperature of the wettest quarter), 9 (mean temperature of the driest quarter), 18 (precipitation of the warmest quarter), and 19 (precipitation of coldest quarter) as they are known to have spatial artifacts that influence model outcomes (Escobar et al. 2014). We also included a variable of bird richness obtained from the IUCN Red List of Threatened Species (IUCN 2023). Three variables related to human activity were included: the human footprint as cumulative human pressure on the environment (Venter et al. 2018), human density, and total human population obtained from the global high resolution population denominators project (WorldPop 2018). Finally, distance of the outbreak to the nearest urban center and to the nearest SAG office were included as proxies of reporting effort, as distance to reporting office can be negatively correlated with probability of reporting (Benavides et al. 2020). We extracted all data for each H5N1 outbreak to coordinates using raster layers of 30 s (∼1 km^2^) spatial resolution with QuantumGIS v.3.8.2104. Table S1 details each assessed variable and data sources.

We built a generalized linear model (GLM) with a binomial error structure (link=logit) to test the association between H5N1 presence/absence model and the variables described above using the glm function in R 4.2.1. As we did not have absence ‘0’ data for the GLM, we extracted variables for pseudo-absence points, randomly selected within a 50 km radius from coordinates of reported outbreaks. Thus, we selected a total of 450 points, equivalent to 2-3 points per outbreak location. A radius of 10 and 100 km generated similar results. We run the full model and tested statistical significance with the Wald’s test. Independent models using similar variables (e.g. three variables of human activities) were tested independently and the model was selected based on their AIC. From the initial WorldClim variables included in the model, we further excluded variable bioclimatic variables 5, 7 and 10 given poor model estimates and/or high correlation with other variables, so these variables are not included in the final presented model. Model fit was evaluated using the *DHARMa* package (functions *testOutliers, testDispersion, qqplot*) in R. The significance of spatial autocorrelation in the residuals of the full model was tested using the Moran’s I test (Glittleman & Kot 1990) in the *spdep* package, and no significant autocorrelation was found.

## RESULTS

### H5N1 occurrence and species affected

A total of 197 outbreaks, involving 478 individual birds tested positive to H5N1 in Chile including 20 domestic chicken (*Gallus gallus*) and 458 wild birds of 26 different species (details in Fig. 1). Most outbreaks involved *P. thagus* (n=72, 37%), followed by Peruvian boobies (*Sula variegata*; 26, 13%), and kelp gulls (*Larus dominicanus*; 20, 10%). The only Endangered species affected was the Humboldt penguin (*Spheniscus humboldti*) presenting 2 outbreaks.

**Figure 1.**
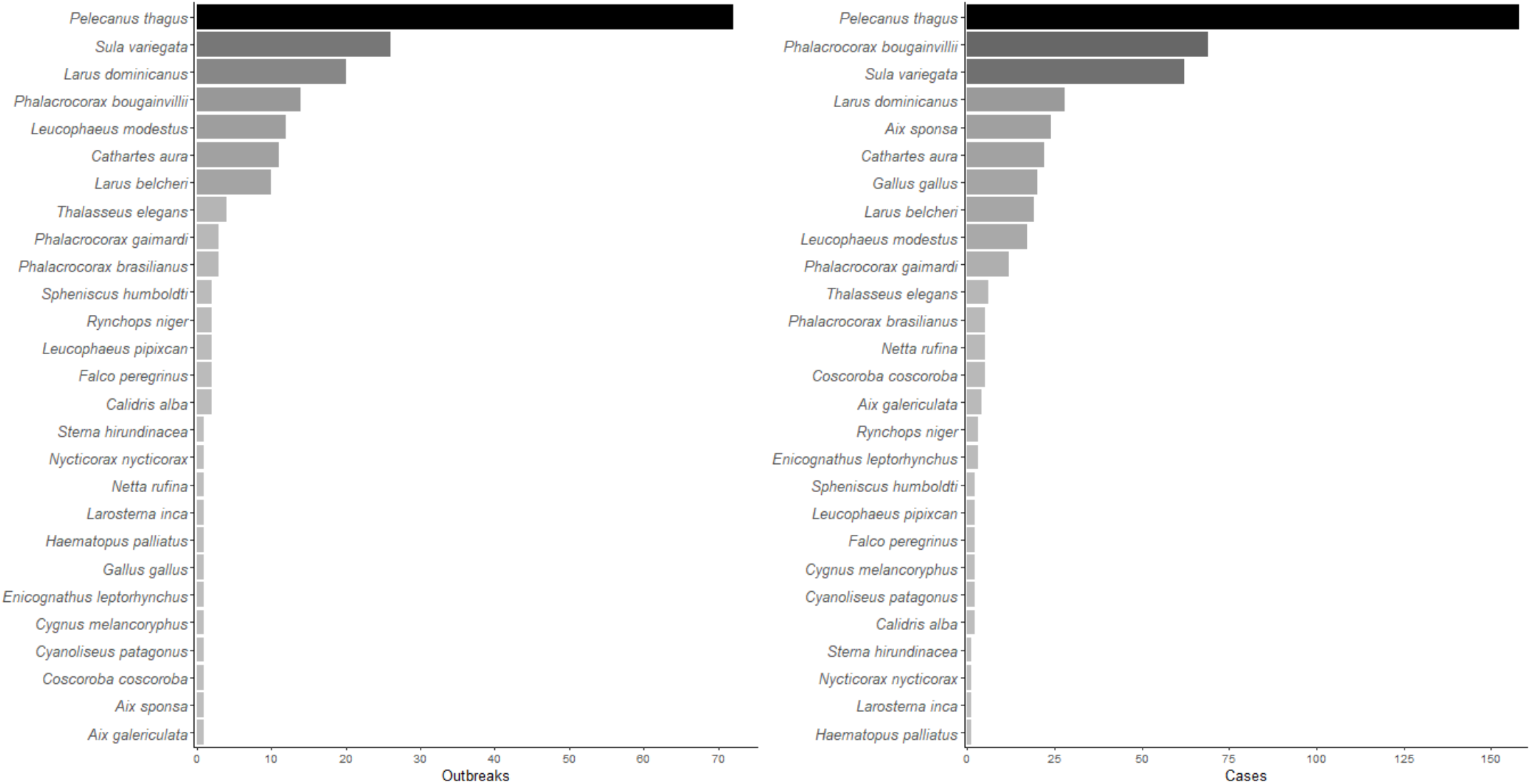
Histogram showing number of a) outbreaks and b) individual cases of highly pathogenic avian influenza H5N1 in birds in Chile between December 2022 and March 2023.

### H5N1 spatio-temporal cluster analysis

Our space-time permutation analysis detected 14 statistically significant clusters of H5N1 outbreaks, following a pattern in which the progression to the south of the H5N1 epidemic wave can be observed (Fig. 2, Table S2). Space-time clusters were scattered from extreme north to south Chile and ranged from two to 19 H5N1 qPCR positives birds in each. Eight of the clusters had a radius of <1 km, while the remaining six clusters were of a variable radius of 4 to 29 km. In addition, four clusters (#1, 2, 7 and 13) were located in the central zone of Chile, close to the most densely populated area in the country. Also, cluster #12 was identified in the northern city of Tocopilla, location where the first H5N1 human case occurred time later (Fig. 2). Chronologically, the epidemic wave of H5N1 was identified from December 2022 in the north (Cluster 9) to February 2023 in the south (Cluster 14). The Global Moran’s I index was statistically non-significant (p=0.2), indicating that there is no spatial autocorrelation of the data.

**Figure 2.**
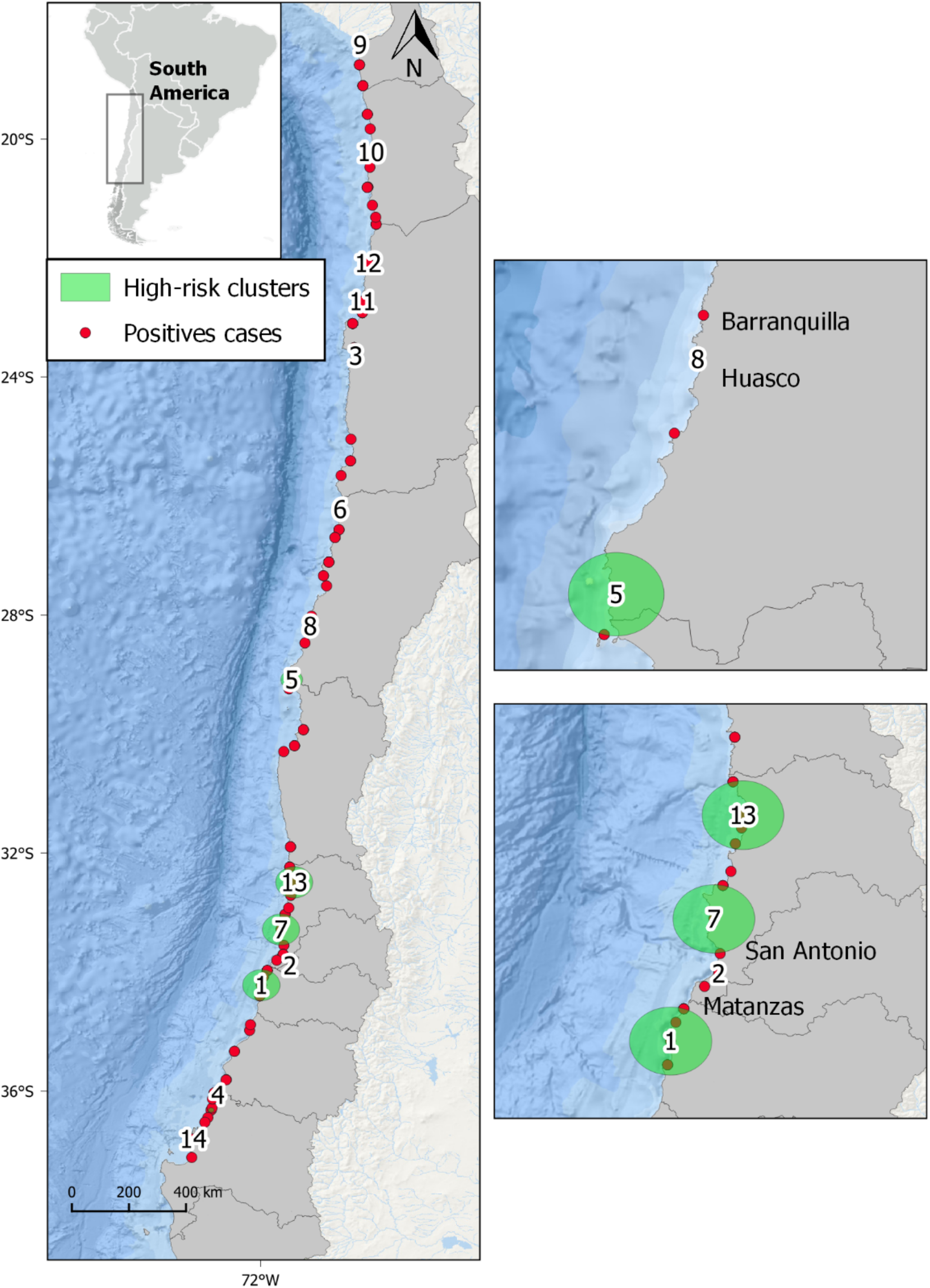
Spatial distribution of highly pathogenic avian influenza H5N1 (H5N1) in birds in Chile studied from December 2022 to February 2023. Distribution of outbreaks, and 14 statistically significant (P<0.05) spatial clusters of H5N1 high-risk areas obtained from scan statistics analysis. Large clusters are represented by green shaded circles.

### Ecological and anthropogenic H5N1 drivers

A strong linear relationship was found between distance and time from the first outbreak (Fig. 3), suggesting a wave-like steady spread. The linear relationship was high (R^2^=0.81) when using grid’s centroids, and substantially improved when excluding four points from Concepción, Cauquenes and Itata municipalities in south Chile (R^2^=0.88).

**Figure 3.**
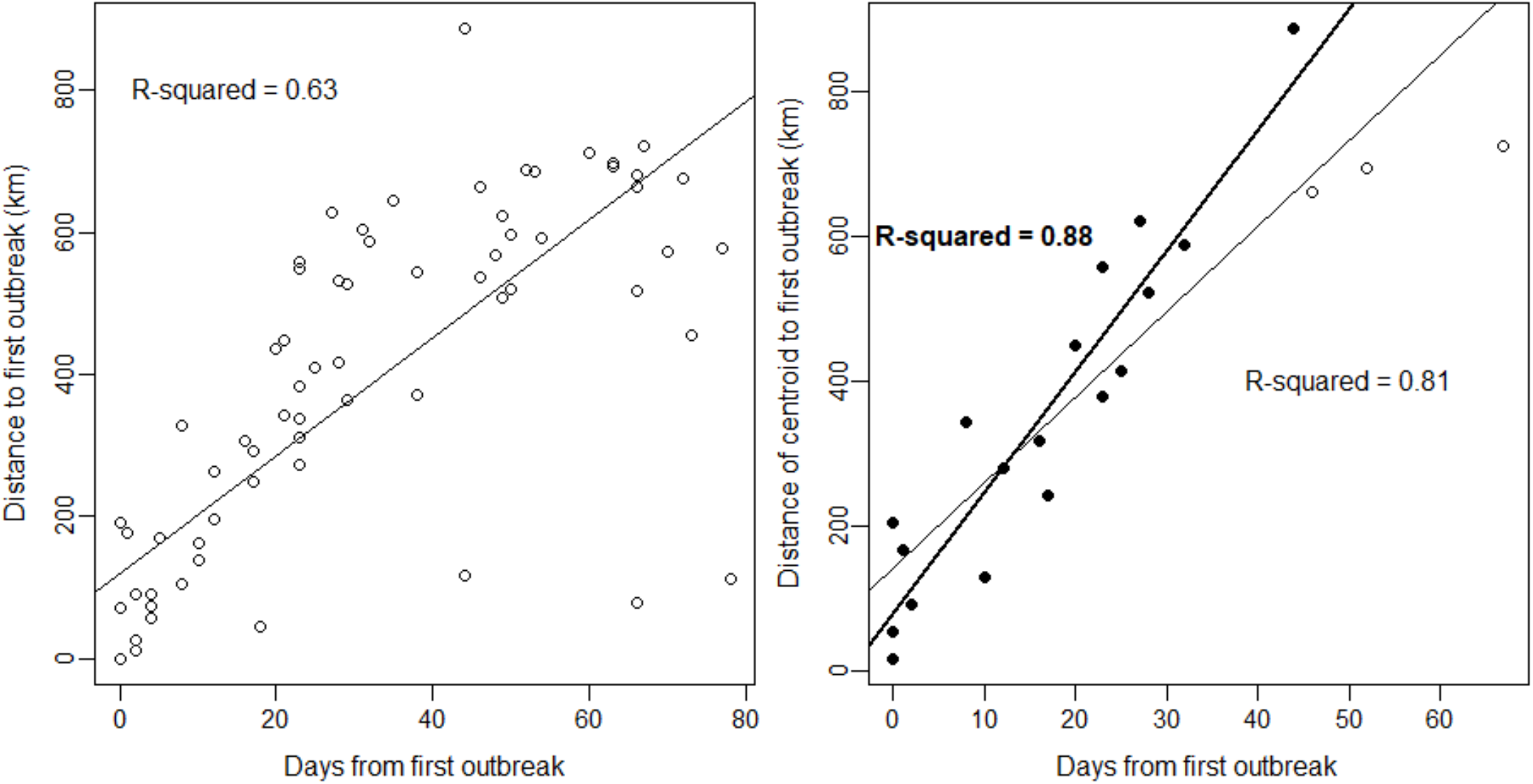
Relationship between distance and days since the first outbreak of highly pathogenic avian influenza H5N1 in birds in Chile. Left: Relationship using coordinates of each outbreak (n=197). The lines are prediction of linear model (R-squared is given for the prediction), with a slope of 8 km/day. Right: Same analysis but using distance from centroids and date of first case detected on the centroid (n= 21), slope of 11.8 km/day. Centroids were estimated using a cell size grid with the *st_make_grid* function of the sf package in R. R-squared and line shown in bold correspond to a linear model excluding points filled in white (n=3) located in the neighboring southern municipalities of Concepcion, Itata and Cauquenes, which substantially improved the model’s R-squared and AIC (from 261 to 214), slope of 16.7 km/day. The first outbreak in the dataset was used as the first index case for all calculations.

The presence of H5N1 outbreaks in birds was correlated to several ecological and human-related variables. Presence of H5N1 was positively correlated to bird richness, human footprint, precipitation of the wettest month, minimum temperature of the coldest month, and mean diurnal range (GLM, p<0.05, Fig. 4, Table S3). In contrast, presence of H5N1 was negatively correlated to distance to the closest urban center, precipitation seasonality and isothermality. No significant association was found between the presence of H5N1 and annual mean temperature, temperature seasonality, mean temperature of the coldest quarter, annual precipitation, precipitation of the driest month, precipitation of the wettest quarter, precipitation of the driest quarter nor distance to the closest SAG office (GLM, p>0.05), nor human total population or density (tested in a separate model including all other variables).

**Figure 4.**
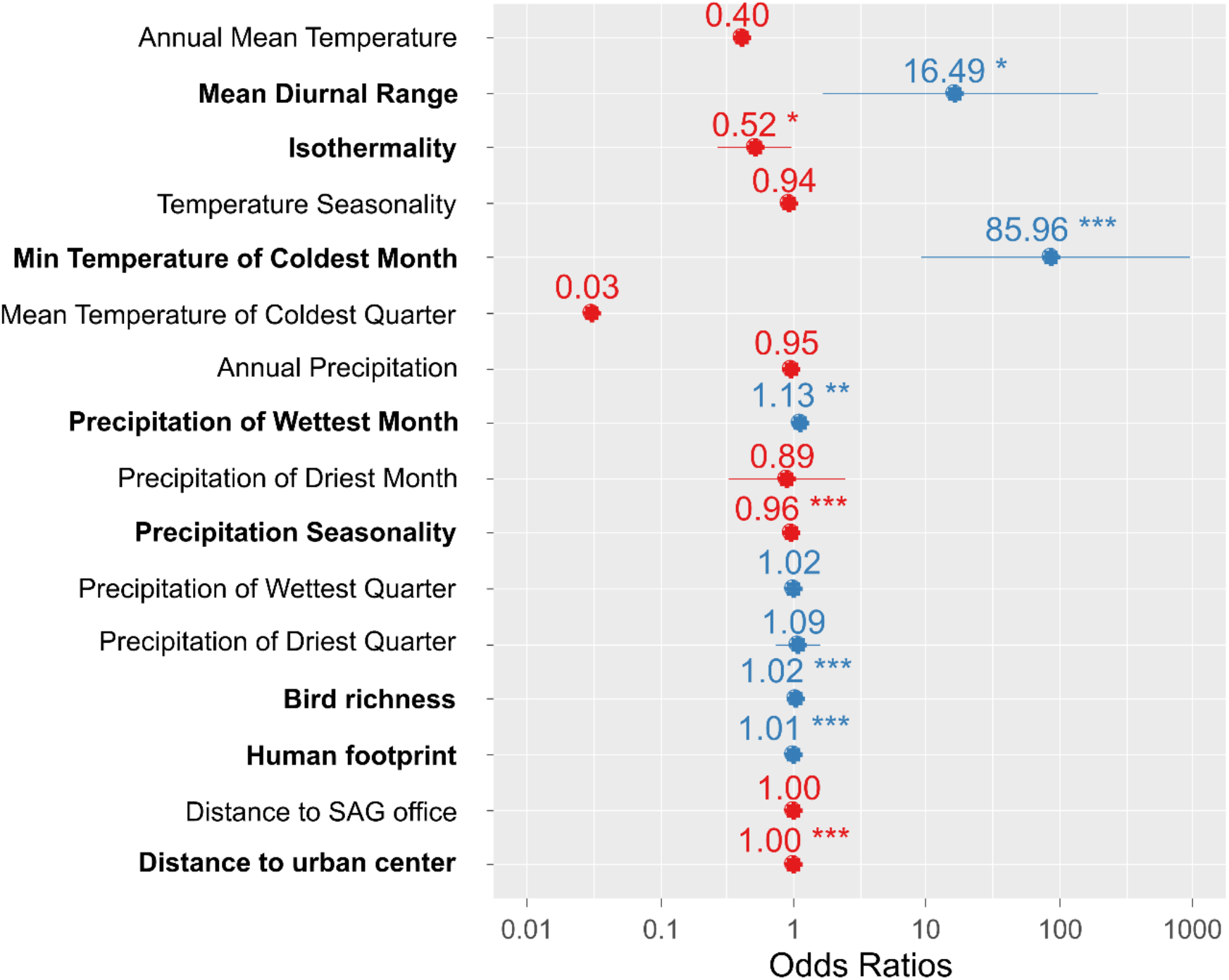
Ecological and anthropogenic variables associated with the presence of highly pathogenic avian influenza H5N1 outbreaks in birds in Chile. Effect size (odds ratio) of variables explaining the presence/absence of cases using a generalized linear model with binomial error distribution. Odds ratios with an asterisk show statistically significant effects (p-value <0.05), the more asterisks, the lower the p-value.

## DISCUSSION

The novel H5N1 virus has rapidly spread worldwide, particularly impacting wildlife and domestic birds across the Americas since late 2021 (Ramey et al. 2022, Gamarra-Toledo et al. 2023a). However, its spatio-temporal dynamics, the epidemiological role of affected hosts, and factors associated with outbreaks remain poorly understood. In this study, we described the dynamics of outbreaks reported in Chile, showing several clusters across the country, a wave-like spread from north to south likely generated by migrated bird arrival during spring, as well as several ecological and human variables associated with outbreaks, from which recommendations for disease management and future research are provided. Our analyses show that the novel H5N1 virus has spread fast across Chile causing high mortality on a diverse wild bird species host range. To date, in four months, the virus has spread 4,000 km along the Pacific flyway in Chile, with infection being reported in 41 wild bird, four domestic bird and four marine mammal species (SAG 2023, SERNAPESCA 2023). Unprecedented mortalities of over 21,000 wild birds and 3,390 marine mammals have been documented in Chile since December 2022 (SAG 2023, SERNAPESCA 2023). This epidemic pattern is similar to the one observed in North America where the virus disseminated fast throughout continental United States and all provinces of Canada, affecting ∼28,000 wild birds among sick and mortality estimates (Harvey et al. 2023), or in Peru, where more than 120,000 wild birds and 3,000 marine mammals have died since October 2022 (Gamarra-Toledo et al. 2023). In addition, in the United States H5N1 has been associated with high mortality rates and breeding failure in the bald eagle (*Haliaeetus leucocephaus*; Nemeth et al. 2023) and unusual mortalities in the highly threatened California condor (*Gymnogyps californianus*) have been recently reported. Likewise, the arrival of H5N1 to Tierra del Fuego in the extreme south of Chile, is now an imminent threat to Antarctic fauna. In continental Chile alone, nearly 1,000 *S. humboldti* have died since January 2023.

Genetic sequencing of Chilean H5N1 samples all cluster together within the clade 2.3.4.4b (Ariyama et al. 2023, Centers for Disease Control and Prevention 2023, Jiménez-Bluhm et al. 2023), suggesting that the outbreaks in the country all belong to the same viral lineage (with potentially minor variations) that is globally widespread, arrived in Chile from neighbor Peru, and previously from the United States (Ramey et al. 2022, Leguia et al. 2023). This wave like spreading is likely associated to the movement of wild birds, although no information is currently available on the hosts species disseminating the virus across space in South America. Although pelicans, boobies, gulls and cormorants have been the most affected species in Chile and Peru (Gamarra-Toledo et al. 2023a), and ducks and teals are recognized as reservoirs and spreaders of avian influenza in other regions (Siembieda et al. 2010, Torrontegi et al. 2019, Ortiz et al. 2023), it is unknown which species are responsible for the observed pattern of spread in Chile, given their movement behavior with most of these species lacking large-scale migrations (Gonzalez-Reiche and Perez 2012, Afanador-Villamizar et al. 2017). Thus, further research combining spatial ecology and genetic analyses could help solving this relevant question.

We showed that outbreak’s occurrence in wild birds and chickens was associated with several bioclimatic variables and proxies of human activities. For example, in our study H5N1 occurrence was negatively associated with distance to urban centers, which could be explained by lower reporting capacity and detection probability as distance from cities increases, despite no significant correlation with distance to the nearest SAG office. The latest absence of correlation could be explained by a larger reporting network beyond these offices, that increases reporting capacity near urban areas (Benavides et al. 2017). For instance, H5N1 qPCR diagnostics is centralized in Santiago (Ariyama et al. 2023) potentially causing a geographic diagnostic bias, which could be further increased by an uneven role of local SAG offices in disease screening. In fact, four of the 14 spatio-temporal clusters of H5N1 outbreaks occurred in central Chile (near Santiago), including the three largest clusters, despite the epizootic followed a north to south spread (Fig. 2). Central Chile also concentrates chicken production in the country. To date, seven outbreaks have been reported in industrial poultry units and 105 in backyard poultry production systems (SAG 2023), from which 3,652 chickens have died and >200,000 have been killed for disease control management (WAHO 2023). Ongoing spillover from wild birds to poultry also threatens Latin America food security, as the region include some of the world largest poultry producers such as Brazil, Mexico, Colombia and Argentina (Food and Agriculture Organization 2021).

Bird species richness and human footprint, but no human density nor total population, were associated with outbreaks in our study, suggesting that areas of high bird diversity and human activities can be hotspots of H5N1, as previously found for bird richness and prevalence of low pathogenic avian influenza viruses in Spanish wetlands (Pérez-Ramírez et al. 2012). Additionally, Huang et al. (2018) found that bird species richness, particularly of higher risk species (i.e., *Larus* spp.), show a positive relationship with H5N1 outbreak probabilities in wild birds in Europe. However, a relationship between bird richness and prevalence of avian influenza was not found previously in wetlands of central Chile (Ruiz et al. 2021). Studies have shown that the black-headed gull (*L. ridibundus*) is highly susceptible to avian influenza in Europe, likely contributing to local transmission rather than long-distance spread due to their high mortality to H5N1 (Ramis et al. 2014). Gulls are among the bird species more involved in outbreaks in Chile (Fig. 1). The largely sedentary behavior and high-density aggregations in colonies during breeding for *L. dominicanus* and Belcher’s gulls (*L. belcheri*) suggest these species may act as competent H5N1 reservoirs contributing with virus persistence in areas such as our identified clusters (Fig. 2). Different is the situation of the grey gull (*L. modestus*), where its migratory condition could help spreading H5N1 along much of the Pacific flyway within South America (Billerman et al. 2023). Conversely, Anseriformes were involved in few outbreaks in Chile (Fig. 1), despite being recognized as key hosts in the epidemiology of avian influenza viruses, both as reservoirs (Gaidet et al. 2012, Verhagen et al. 2014, Jiménez-Bluhm et al. 2018, Torrontegi et al. 2019, Ortiz et al. 2023) or as long-distance spreaders (Keawcharoen et al. 2008). As previously shown for a wide variety of low pathogenic avian influenza viruses in Chile (Jiménez-Bluhm et al. 2018, Ruiz et al. 2021), it is possible that as H5N1 becomes enzootic in Chile, Anseriformes start playing more relevant epidemiological roles, such as acting as virus reservoirs (Torrontegi et al. 2019, Pohlmann et al. 2022) or even suffering the consequences of disease (Keawcharoen et al. 2008). For instance, the unusual mortality of over 500 black-necked swans (*C. melanocoryphus*) in a wetland in Valdivia (southern Chile) associated with H5N1 is of high concern (SAG 2023).

Among climatic variables, the minimum temperature of the coldest month (∼86 times more likely per temperature degree) and mean diurnal range (∼16 times per temperature degree) has the highest positively association with outbreaks, encouraging research on ecological modeling predictions of future outbreaks, particularly for the upcoming critical winter period, which have shown a peak of outbreaks of H5N1 in North America (Harvey et al. 2023) and Europe (Pohlmann et al. 2022, European Food safety Authority et al. 2023). Similarly, in a study in central Chilean wetlands and based on fecal samples of wild birds, Ruiz et al. (2021) found a positive association of prevalence of low pathogenic avian influenza viruses with minimum temperature for the month of sampling. But contrasting results were obtained for the prevalence of avian influenza viruses in feces of wild birds in Spain, where monthly mean lowest daily temperature was negatively correlated with prevalence of avian influenza (Pérez-Ramírez et al. 2012). We did not find a significant correlation between outbreak occurrence and annual precipitation, as also reported by Ruiz et al. (2021). Although previous studies have found a higher prevalence of enzootic avian influenza viruses during summer and autumn in Chile (Jiménez-Bluhm et al. 2018, Ruiz et al. 2021), the progression of the current H5N1 epizootic in South America will allow to test for seasonality effects. Also, as the epizootic moves inland, it is important to include in future studies the analysis of other ecological variables that might be relevant, such as vegetation coverage and the size of water bodies (Ruiz et al. 2021) and host density (Gaidet et al. 2012). The biological mechanisms behind the observed associations found in our study with several bioclimatic variables (e.g., precipitation seasonality or isothermatlity) in terms of host and viral dynamics should be matter of further research.

One of the most worrying issues of the current panzootic is the increasing cases of H5N1 infections in wild mammals, both marine and terrestrial, causing unprecedented mass mortalities (Ramey et al. 2022, Bordes et al. 2023, Gamarra-Toledo et al. 2023b, Leguia et al. 2023, Puryear et al. 2023). Worryingly, RNA sequencing has shown signs of early adaptation of the H5N1 2.3.4.4b clade virus to wild mammals (Vreman et al. 2023) and humans (Centers for Disease Control and Prevention 2023), fueling the fears of a new human pandemic (Kupferschmidt 2023). In addition, recent reports of a dog and two cats dying from H5N1 after contact with dead birds in North America (Canadian Food Inspection Agency 2023) rises the potential of wildlife to dog transmission in South American countries like Chile, where a high population of dogs (both owned and stray) occurs on the coast (Cortés et al. 2021). Even more, two cases of zoonotic avian influenza have recently been reported in South America (Ecuador and Chile), with the Chilean case being of special concern. Here, the location of this case (Tocopilla) was predicted as a high-risk area by our spatio-temporal permutation analysis (cluster #12, Fig. 2). The epidemiological investigation concluded that the transmission most likely occurred through environmental exposure due to the high amount of dead sea lions and wild birds near the infected man’s residence (Centers for Disease Control and Prevention 2023, World Health Organization 2023), showing the potential of H5N1 spillover to humans from wildlife even in the absence of direct contact.

Since its original description in southern China in 1996 (Wan 1998), H5N1 is causing its highest impacts and has become more widespread than ever (Ramey et al. 2022, Shi et al. 2023). Our results identified a wave-like steady spreading of the novel H5N1 virus in Chile, predicted hotspots of H5N1 risk and factors associated to outbreaks occurrence, contributing to establish targeted preventive measures to avoid further spread and spillover to other species, including humans. For example, regular wave-like patterns of vampire bat rabies in Peru have contributed to alert communities ahead of the spreading wave to implement preventive strategies such as vaccination (Benavides et al. 2016). In our study, potential wave slow-down or late reporting in the area surrounded Concepción might explain divergence with a faster and more steady spread of around 16.7 km/day (Fig. 3). Traditionally, strategies to control avian influenza outbreaks have focused on poultry (Sidik 2023, Shi et al. 2023), but new and urgent efforts are needed to contain viral circulation in wild populations and the environment (Ramey et al. 2022). Preventive actions based on our modeling approach include developing wildlife surveillance diagnostic capabilities in Chilean regions concentrating outbreaks. It is also needed that scientists, local communities, the poultry sector and national health authorities co-design and implement science-based measures from a One Health perspective to avoid further spillover to domestic animals and humans, including rapid removal and proper disposal of wild dead animals, and the closure of public areas (i.e., beaches) reporting high wildlife mortalities. Likewise, target surveillance of key wild species including bird banding, wetland monitoring and citizen science could all contribute to a better understanding of the epidemiology of H5N1. Increasing wildlife disease surveillance and genomic sequencing is also essential to better understand cross-species transmission rates and how the virus is potentially adapting to mammals. Our work highlights the urgent need for further research to identify, alert and prevent hotspots of H5N1 transmission, limiting the potential for a new pandemic emergence, food security catastrophe and further biodiversity loss.

## Acknowledgements

We are grateful of Andrew M. Ramey for his critical review of an early version which help substantially to improve the manuscript.

## SUPPLEMENTARY MATERIAL

**Table S1.** Explanatory variables used in the generalized linear model of highly pathogenic avian influenza H5N1 in Chile.

| Variable | Description | Source |
| --- | --- | --- |
| Bioclimate | 15 bioclimatic variables representative of the period 1970-2000 with a resolution of 30-arc seconds (~1 km) from https://www.worldclim.org including: 1) annual mean temperature, 2) mean diurnal range, 3) isothermality, 4) temperature seasonality, 5) maximum temperature of warmest month, 6) minimum temperature of coldest month, 7) temperature annual range, 10) mean temperature of warmest quarter, 11) mean temperature of coldest quarter, 12) annual precipitation, 13) precipitation of wettest month, 14) precipitation of driest month, 15) precipitation seasonality, 16) precipitation of wettest quarter, 17) precipitation of driest quarter. Variables 8, 9, 18 and 19 were not included. | Fick & Hijmans et al. 2017 |
| Bird richness | Two variables representing bird richness at a spatial resolution of ~1 km, obtained by overlaying the distribution range maps of the IUCN Red List of threatened species from https://www.iucnredlist.org. One included all species, and the second included threatened species. | IUCN 2023 |
| Human Footprint | Cumulative human pressure on the environment at a spatial resolution of ~1 km obtained from https://sedac.ciesin.columbia.edu/data/set/wildareas-v3-2009-human-footprint. The human pressure is measured using eight variables including built-up environments, population density, electric power infrastructure, crop lands, pasture lands, roads, railways, and navigable waterways. | Venter et al. 2018 |
| Human population | Two variables at a spatial resolution of ~1 km obtained from https://www.worldpop.org: 1) estimated population density, and 2) estimated number of people. | WorldPop 2018 |
| Distance to the nearest urban center | Distance in meters from outbreaks points to the centroids of populated areas polygon shapefile, obtained from https://www.bcn.cl/siit/mapas_vectoriales. Spatial data projected in WGS 84 / UTM zone 19S coordinate reference system. | Geospatial Data Infrastructure of Chile 2023 |
| Distance to nearest SAG office | Distance in meters from the outbreak to the regional SAG office location. SAG office geographical coordinates were downloaded from the web map service https://geoportal.sag.gob.cl of the SAG's Department of Geospatial Information Systems. Spatial data projected in WGS 84 / UTM zone 19S coordinate reference system. | Geoportal SAG 2023 |

**Table S2.** Statistically significant clusters (P<0.05) detected by spatiotemporal permutation model using the space–time scan statistic for the novel highly pathogenic avian influenza H5N1 outbreaks between December 2022 and February 2023 in Chile. S: south; W: west; O: observed outbreaks; E: expected outbreaks; and O/E: the observed/expected ratio.

| Cluster | Region | Latitude | Longitude | Radius (km) | Time frame | O | E | O/E | P-value |
| --- | --- | --- | --- | --- | --- | --- | --- | --- | --- |
| 1 | O'Higgins | -34.21 S | -71.98 W | 28.9 | 2023/1/4 to 2023/1/10 | 11 | 0.99 | 11.11 | < 0.001 |
| 2 | Valparaíso | -33.7 S | -71.61 W | 0 | 2023/2/8 to 2023/2/14 | 7 | 0.43 | 16.42 | < 0.001 |
| 3 | Antofagasta | -23.64 S | -70.39 W | 0 | 2022/12/14 to 2022/12/20 | 6 | 0.52 | 11.59 | < 0.001 |
| 4 | Biobio | -36.31 S | -72.82 W | 4.27 | 2023/1/25 to 2023/1/31 | 3 | 0.06 | 49.25 | < 0.001 |
| 5 | Atacama | -29.08 S | -71.47 W | 17.74 | 2022/12/28 to 2023/1/3 | 6 | 0.67 | 8.95 | 0.002 |
| 6 | Atacama | -26.22 S | -70.66 W | 0 | 2022/12/21 to 2022/12/27 | 5 | 0.53 | 9.38 | 0.008 |
| 7 | Valparaíso | -33.28 S | -71.65 W | 29.05 | 2022/12/28 to 2023/1/3 | 5 | 0.56 | 8.95 | 0.011 |
| 8 | Atacama | -28.18 S | -71.16 W | 0 | 2023/1/11 to 2023/1/17 | 2 | 0.03 | 65.67 | 0.010 |
| 9 | Arica | -18.41 S | -70.32 W | 0 | 2022/12/1 to 2022/12/13 | 19 | 7.62 | 2.49 | 0.011 |
| 10 | Tarapacá | -20.21 S | -70.14 W | 13.26 | 2022/12/1 to 2022/12/13 | 19 | 7.62 | 2.49 | 0.011 |
| 11 | Antofagasta | -22.72 S | -70.28 W | 0 | 2022/12/14 to 2022/12/20 | 4 | 0.35 | 11.59 | 0.013 |
| 12 | Antofagasta | -22.08 S | -70.19 W | 0 | 2022/12/14 to 2022/12/20 | 4 | 0.35 | 11.59 | 0.013 |
| 13 | Valparaíso | -32.49 S | -71.43 W | 29.24 | 2023/1/18 to 2023/1/24 | 5 | 0.64 | 7.82 | 0.014 |
| 14 | Biobio | -36.8 S | -73.12 W | 0 | 2023/2/1 to 2023/2/7 | 2 | 0.04 | 49.25 | 0.028 |

**Table S3.**
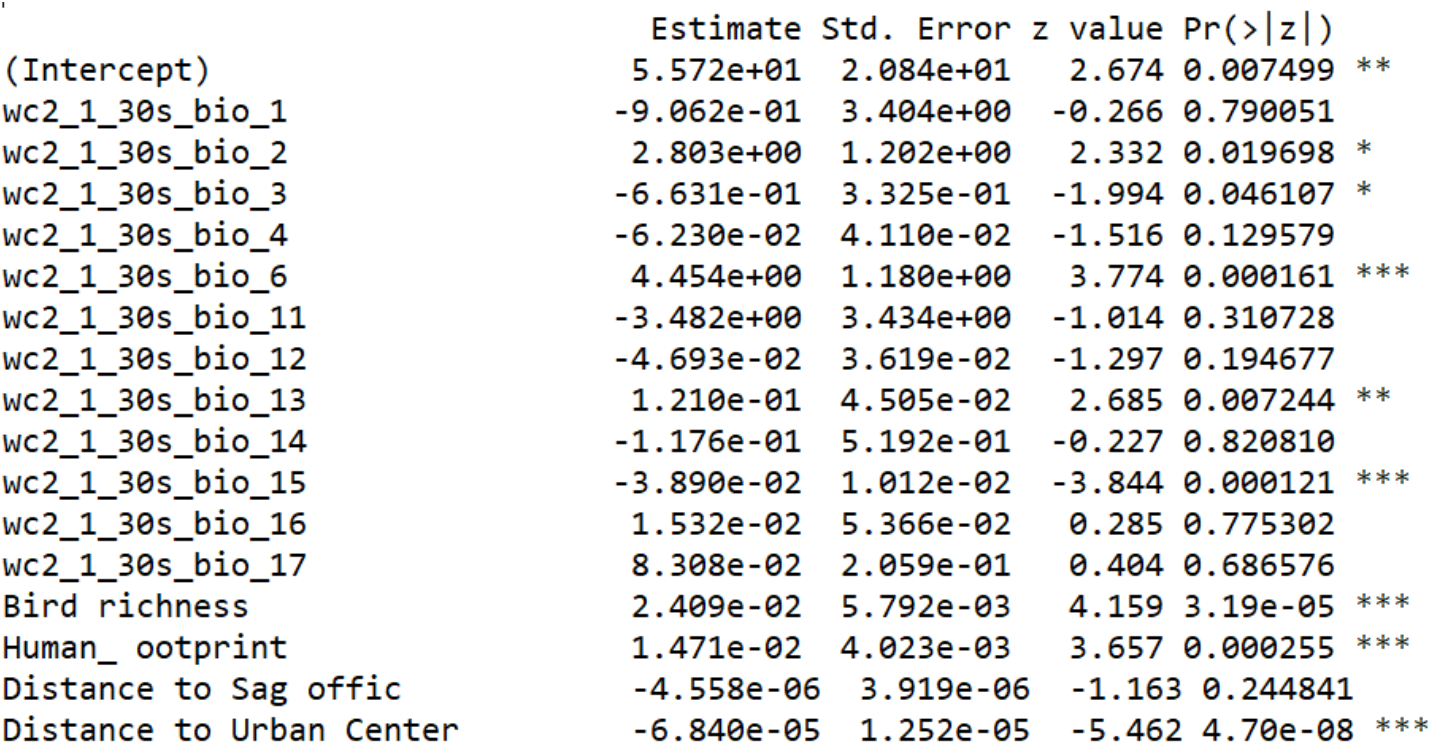
Result of the general lineal model with Binomial residual error explaining highly pathogenic avian influenza H5N1 outbreak presence in Chile.

|  | Estimate | Std. Error | z value | Pr(>\|z\|) |
| --- | --- | --- | --- | --- |
| (Intercept) | 5.572e+01 | 2.084e+01 | 2.674 | 0.007499 ** |
| wc2_1_30s_bio_1 | -9.062e-01 | 3.404e+00 | -0.266 | 0.790051 |
| wc2_1_30s_bio_2 | 2.803e+00 | 1.202e+00 | 2.332 | 0.019698 * |
| wc2_1_30s_bio_3 | -6.631e-01 | 3.325e-01 | -1.994 | 0.046107 * |
| wc2_1_30s_bio_4 | -6.230e-02 | 4.110e-02 | -1.516 | 0.129579 |
| wc2_1_30s_bio_6 | 4.454e+00 | 1.180e+00 | 3.774 | 0.000161 *** |
| wc2_1_30s_bio_11 | -3.482e+00 | 3.434e+00 | -1.014 | 0.310728 |
| wc2_1_30s_bio_12 | -4.693e-02 | 3.619e-02 | -1.297 | 0.194677 |
| wc2_1_30s_bio_13 | 1.210e-01 | 4.505e-02 | 2.685 | 0.007244 ** |
| wc2_1_30s_bio_14 | -1.176e-01 | 5.192e-01 | -0.227 | 0.820810 |
| wc2_1_30s_bio_15 | -3.890e-02 | 1.012e-02 | -3.844 | 0.000121 *** |
| wc2_1_30s_bio_16 | 1.532e-02 | 5.366e-02 | 0.285 | 0.775302 |
| wc2_1_30s_bio_17 | 8.308e-02 | 2.059e-01 | 0.404 | 0.686576 |
| Bird richness | 2.409e-02 | 5.792e-03 | 4.159 | 3.19e-05 *** |
| Human_ ootprint | 1.471e-02 | 4.023e-03 | 3.657 | 0.000255 *** |
| Distance to Sag offic | -4.558e-06 | 3.919e-06 | -1.163 | 0.244841 |
| Distance to Urban Center | -6.840e-05 | 1.252e-05 | -5.462 | 4.70e-08 *** |

